# PyADM1: a Python implementation of Anaerobic Digestion Model No. 1

**DOI:** 10.1101/2021.03.03.433746

**Authors:** P. Sadrimajd, P. Mannion, E. Howley, P. N. L. Lens

**Author notes:** Email addresses* (P. Sadrimajd), (P. Mannion), (E. Howley), (P. N. L. Lens).

## Abstract

Anaerobic Digestion (AD) is a waste treatment technology widely used for wastewater and solid waste treatment, with the advantage of being a source of renewable energy in the form of biogas. Anaerobic digestion model number 1 (ADM1) is the most common mathematical model available for AD modelling. Commercial software implementations of ADM1 are available but have limited flexibility and availability due to the closed sources and licensing fees. Python is the fastest growing programming language and is open source freely available. Python implementation of ADM1 makes this AD model available to the mass user base of the Python ecosystem and it’s libraries. The open easy to use implementation in PyADM1 makes it more accessible and provides possibilities for flexible direct use of the model linked to other software, e.g. machine learning libraries or Linux operating system on embedded hardware.

## Required Metadata

### Current code version

**Table 1:** Code metadata

| Nr. | Code metadata description | Please fill in this column |
| --- | --- | --- |
| C1 | Current code version | 0.3 |
| C2 | Permanent link to code/repository used for this code version | https://github.com/CaptainFerMag/PyADM1 |
| C3 | Code Ocean compute capsule | https://codeocean.com/capsule/0554841/tree |
| C4 | Legal Code License | MIT |
| C5 | Code versioning system used | git |
| C6 | Software code languages, tools, and services used | Compilation not required; Windows, Mac OSX, Linux, Python, SciPy, NumPy |
| C7 | Compilation requirements, operating environments & dependencies | Copy, Scipy, Numpy, Python 3.8 |
| C8 | If available Link to developer documentation/manual | https://github.com/CaptainFerMag/PyADM1 |
| C9 | Support email for questions | p.sadrimajd1[at]nuigalway.ie |

## 1. Motivation and significance

Large amounts of organic waste (e.g. food waste, municipal wastewater, industrial wastewater, and agricultural waste) are being disposed every year. Compared to traditional waste treatment (i.e. landfill and composting) Anaerobic Digestion (AD) not only reduces emissions to the environment, but also produces renewable energy in the form of biogas [1]. There are numerous biogas plants around the world and with the growing concerns about climate change there will be more biogas plants in the future. According to the International Energy Agency (IEA), in 2019 there were 10000 biogas plants in Germany, 1000 in UK, and 400 in Switzerland [2].

Anaerobic Digestion Model number 1 (ADM1) from the International Water Association (IWA) [3] (the most commonly used model for AD) describes AD as a five steps process (disintegration, hydrolysis, acidogenesis or fermentation, acetogenesis, and methanogenesis). ADM1 is a complex mathematical model that includes 35 state variables and their respective differential or algebraic equations describing the interrelated change between states [4]. This makes the use of computers and numerical solvers necessary for ADM1 application.

ADM1 has been implemented in different commercial softwares (e.g. GPS-X, SIMBA, WEST, MATLAB/SIMULINK, and AQUASIM) and programming languages (i.e. JAVA, FORTRAN, and C) [5, 6, 7, 8]. However, commercial packages involve license costs and are not flexible (configurable) enough for automation of modeling workflows and applications, and for model customization and updates. Benchmark Simulation Model number 2 (BSM2) [6] developed in the MATLAB/SIMULINK environment is one of the most widely used open source ADM1 implementations. BSM2 is available in the MATLAB/SIMULINK environment where the model code is available in C programming language. BSM2 was developed in older versions of MATLAB and requires modifications to run properly in MATLAB versions after 2017. In order to use BSM2 a purchase of MATLAB license is required and there is a learning curve involved in using MATLAB/SIMULINK environment.

Python being a freely available open source programming language, is the fastest growing programming language and the first programming language that many young scientists learn [9]. Also, many modern scientific programming and data analysis libraries (e.g. SciPy [10]) are developed in Python. Implementing ADM1 in Python makes it more accessible and easier to use for a wider community of researchers and engineers, enabling them to expand ADM1 applications and anaerobic digestion research.

## 2. Software description

PyADM1 is a Python implementation of ADM1 based on BSM2 that can be used with basic Python programming knowledge in any platform running Python. PyADM1 uses `scipy.integrate.solve_ivp` to solve systems of differential equations and `numpy` for math operations.

### 2.1. Software Architecture

The general Ordinary Differential Equation (ODE) form of ADM1 is:

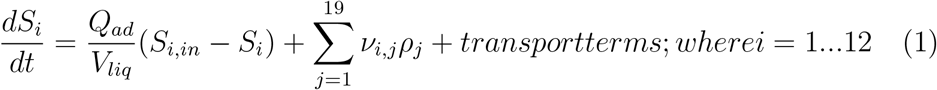

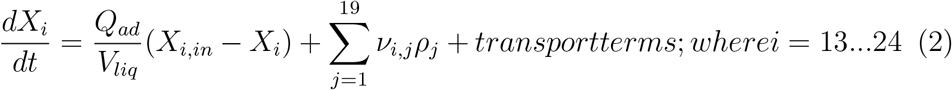

where *Q_ad_* is influent flow rate, *V_liq_* is the volume of the liquid fraction of reactor, *S_i_* and *X_i_* are solubles matter and particular matter states for variable *i*, respectively. The terms *ν* (Biochemical rate coefficients) and *ρ* (kinetic equations for process *j*) are from the Peterson Matrix format of ADM1 [3]. Equations 25 and 26 are related to cations and anions, respectively. Equations 27-32 are for ion states and differ between the ODE and Differential Algebraic Equations (DAE) form of ADM1. Equations 33-35 are related to the gas phase.

The software architecture is presented in Figure 1. Initial states (*S_i_* and *X_i_*), influent states (*S_i,in_* and *X_i,in_*), simulation time (`days`), number of time steps (`timeSteps`), and model parameters are defined as global variables which are modifiable by the user, depending on the simulation scenario or experimental data [3, 11, 12, 13] for modeling purposes. After integration and solving the system of model equations over the given time period, PyADM1 delivers the results of the simulation as system state outputs (*S_i_*). The current version of PyADM1 is capable of steady-state simulations with DAE and ODE forms of ADM1. Dynamic simulation capabilities will be added to the future versions of PyADM1. A user can switch between ODE/DAE modes by setting the `DAE_switch` variable to 0 for ODE and 1 for DAE.

**Figure 1:**
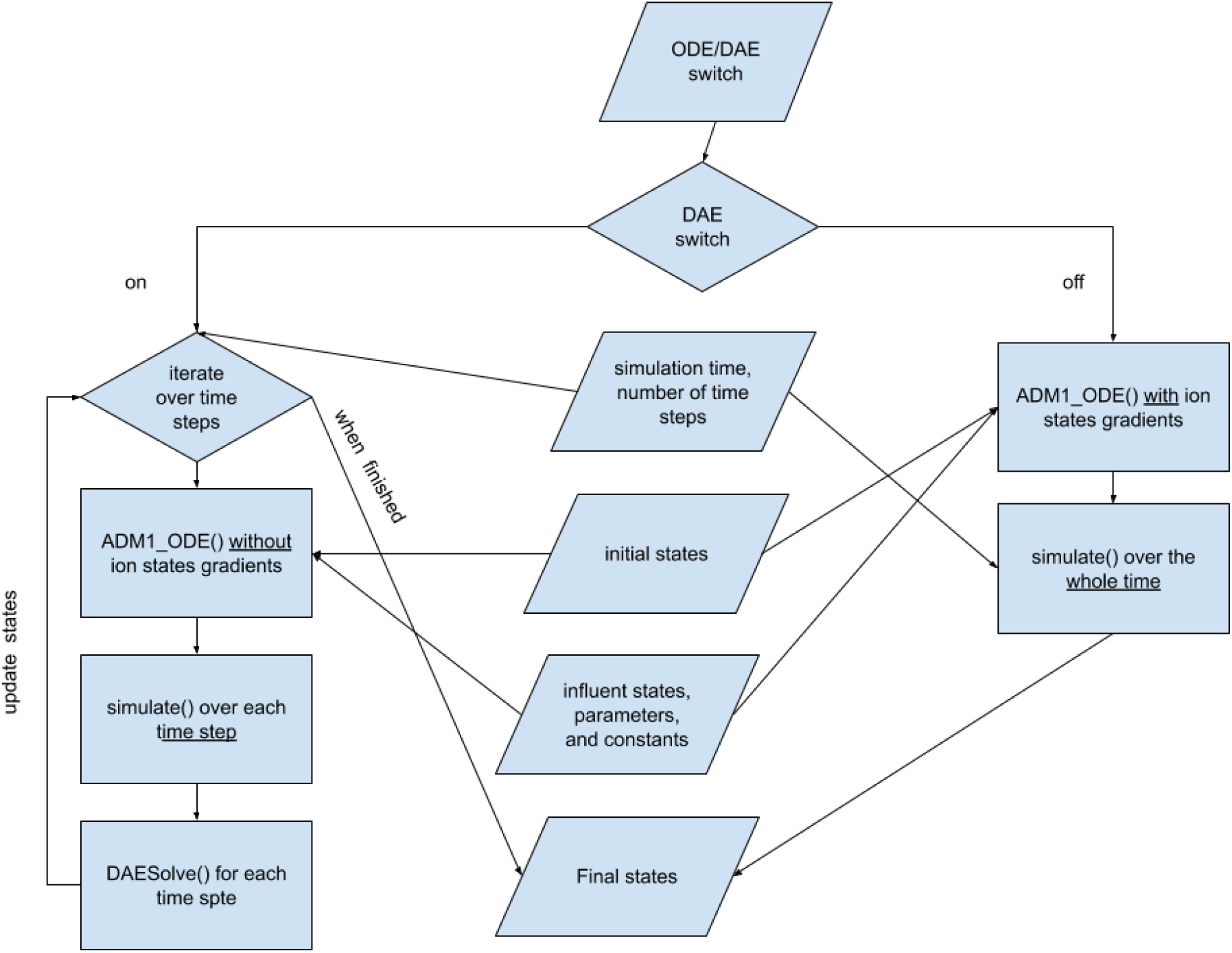
PyADM1 code architecture and flow of calculations

### 2.2. Software Functionalities

There are three main functions. Function `ADM1_ODE(t, state_zero)` receives simulation time *t* and initial states *state zero* as input and calculates the derivatives of states as an ODE. Function `simulate(t_step, solvermethod)` uses `scipy.integrate.solve_ivp()` to solve the initial value problem (ivp) integration at each time step and integrate the ODE equations. In other words, it receives the initial states at the beginning of each time step and solves the differential equations to deliver the state values at the end of the time step. Function DAESolve() calculates the derivatives for 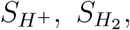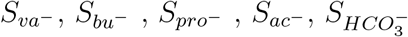, and 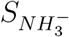.

Variable `DAE_switch` is to choose between complete ODE form (`DAE_switch=0`) and DAE form (`DAE_switch=1`) of ADM1 for the simulation. If `DAE_switch=0` then `ADM1_ODE()` includes ion state gradients in the ODE system and `simulate()` integration is over the whole simulations time and the final states in the end of the simulation time are reported. Hence, if `DAE_switch=1` then a loop iterates over time steps and at each time step `ADM1_ODE()` excludes ion states from the ODE and `simulate()` integration is only over the current time step. Further, the simulated states in the current time step are delivered to `DAESolve()` to calculate the rest of the states at the end of the time step. Then the iteration over the current state is over and the loop moves to the next time step until the end of the simulation time.

## 3. Illustrative Examples

In order to validate the PyADM1 code functionality, a steady-state simulation was executed using the exact same input states, initial states, and parameters as the BSM2 (default values for state and parameter variables in the PyADM1 code). Simulation time was 200 days divided into 15 minutes intervals (i.e. time steps) and substrate characterisation (defines influent states) was based on wastewater treatment sludge.

PyADM1 DAE steady-state simulation inputs, results, and comparative errors are presented in Tables 2, 3 and 4. Decimal places reported here are based on the BSM2 report for the purpose of demonstrating numerical calculations accuracy and the significance should be based on the unit of measurement for each state (commented in front of each state in the source code). Achieved absolute error levels for all simulated states were smaller than 10^−5^ (i.e. it has no physical significance in relation to the units of measurement for process states) and according to the BSM2 report [6] this error level validates the PyADM1 model implementation.

**Table 2:** Steady state simulation results comparison between BSM2 DAE2 Matlab implementation and PyADM1 DAE (soluble matter)

| state | Influent | Initial | BSM2 | PyADM1 | Error |
| --- | --- | --- | --- | --- | --- |
| $S_{su}$ | 0.01 | 0.012 | 0.01195482971696 | 0.01195482971230 | 5E-12 |
| $S_{aa}$ | 0.001 | 0.0053 | 0.00531474017163 | 0.00531474016819 | 3E-12 |
| $S_{fa}$ | 0.001 | 0.099 | 0.09862140093080 | 0.09862140187903 | -9E-10 |
| $S_{va}$ | 0.001 | 0.012 | 0.01162500646386 | 0.01162500650562 | -4E-11 |
| $S_{bu}$ | 0.001 | 0.013 | 0.01325072966627 | 0.01325072971277 | -5E-11 |
| $S_{pro}$ | 0.001 | 0.016 | 0.01578366628454 | 0.01578366597247 | 3E-10 |
| $S_{ac}$ | 0.001 | 0.2 | 0.19762971693755 | 0.19763167669097 | -2E-06 |
| $S_{H_2}$ | 10E-8 | 2.30E-7 | 0.00000023594510 | 0.00000023594506 | 4E-14 |
| $S_{CH_4}$ | 10E-5 | 0.055 | 0.05508877644596 | 0.05508904965331 | -3E-07 |
| $S_{IC}$ | 0.04 | 0.15 | 0.15267787062633 | 0.15267747569836 | 4E-07 |
| $S_{IN}$ | 0.01 | 0.13 | 0.13022981580368 | 0.13022980517356 | 1E-08 |
| $S_I$ | 0.02 | 0.33 | 0.32869766372153 | 0.32869772786660 | -6E-08 |

**Table 3:** steady state simulation results comparison between BSM2 DAE2 Matlab implementation and PyADM1 DAE (particulate matter)

| state | Influent | Initial | BSM2 | PyADM1 | Error |
| --- | --- | --- | --- | --- | --- |
| $X_{xc}$ | 2 | 0.31 | 0.30869766372153 | 0.30869765844290 | 5E-09 |
| $X_{ch}$ | 5 | 0.028 | 0.02794724043504 | 0.02794724038020 | 5E-11 |
| $X_{pr}$ | 20 | 0.1 | 0.10257410610668 | 0.10257410605184 | 5E-11 |
| $X_{li}$ | 5 | 0.029 | 0.02948304970729 | 0.02948304962503 | 8E-11 |
| $X_{su}$ | 0.0 | 0.42 | 0.42016598245457 | 0.42016598174665 | 7E-10 |
| $X_{aa}$ | 0.01 | 1.18 | 1.17917179892370 | 1.17917179900952 | -9E-11 |
| $X_{fa}$ | 0.01 | 0.24 | 0.24303534471944 | 0.24303534195974 | 3E-09 |
| $X_{C4}$ | 0.01 | 0.43 | 0.43192110563598 | 0.43192110381834 | 2E-09 |
| $X_{pro}$ | 0.01 | 0.14 | 0.13730590893395 | 0.13730591105219 | -2E-09 |
| $X_{ac}$ | 0.01 | 0.76 | 0.76056265831322 | 0.76056260560337 | 5E-08 |
| $X_{H_2}$ | 0.01 | 0.32 | 0.31702295336127 | 0.31702295557648 | -2E-09 |
| $X_I$ | 25.0 | 25.6 | 25.61739532744300 | 25.61739454773450 | 8E-07 |

**Table 4:** steady state simulation results comparison between BSM2 DAE2 Matlab implementation and PyADM1 DAE (ion and gas)

| state | Influent | Initial | BSM2 | PyADM1 | Error |
| --- | --- | --- | --- | --- | --- |
| $S_{cation}$ | 0.04 | 0.040 | 0.04 | 0.04 | 0E+00 |
| $S_{anion}$ | 0.02 | 0.020 | 0.02 | 0.02 | 0E+00 |
| $S_{H^+}$ | 1.0 | 0.00000003423 | 0.00000003423440 | 0.00000003423351 | 9E-13 |
| $S_{va^-}$ | 0.0 | 0.011 | 0.01159624707260 | 0.01159624782654 | -8E-10 |
| $S_{bu^-}$ | 0.0 | 0.013 | 0.01322082624850 | 0.01322082703576 | -8E-10 |
| $S_{pro^-}$ | 0.0 | 0.016 | 0.01574278319160 | 0.01574278389282 | -7E-10 |
| $S_{ac^-}$ | 0.0 | 0.2 | 0.19724115543650 | 0.19724312096623 | -2E-06 |
| $S_{HCO_3^-}$ | 0.0 | 0.14 | 0.14277747939210 | 0.14277733996334 | 1E-07 |
| $S_{CO_2}$ | 0.0 | 0.0 | 0.00990039123430 | 0.00990013573502 | 3E-07 |
| $S_{NH_3}$ | 0.0 | 0.0041 | 0.00409092845840 | 0.00409102651559 | -1E-07 |
| $S_{NH_4^+}$ | 0.0 | 0.0 | 0.12613888734520 | 0.12613877865798 | 1E-07 |
| $S_{gas,H_2}$ | 0.0 | 1.02E-5 | 0.00001024103560 | 0.00001024110692 | -7E-11 |
| $S_{gas,CH_4}$ | 0.0 | 1.63 | 1.62560720998142 | 1.62562665039951 | -2E-05 |
| $S_{gas,CO_2}$ | 0.0 | 0.014 | 0.01415053467840 | 0.01415239461914 | -2E-06 |

## 4. Impact

There are about 30000 peer-reviewed entries related to “Anaerobic Digestion” search term available in the Scopus database. Establishing the correctness of what is published (i.e. benchmarking approach) a Python implementation of ADM1 will make it available for many more researchers and opens new possibilities for extension and configuration of ADM1. It will allow researchers to reproduce published work and compare the results of the experiments in different scenarios and for different digester regimes, e.g. batch or semi-batch, continuous, up flow, and circulation.

In contrast to commercial packages, the open source code of PyADM1 facilitates model modifications by the user in a modern programming language. More than 6000 search results show up on Google Scholar for the query “ADM1 modified anaerobic digestion” results. Based on the niche application, model modifications can add equations to simulate additional states (i.e. nutrients, nitrogen, phosphorus, sulphur, and metals), introduce the effect of temperature on the model parameters, or different microbial groups (as new types of particulate matter). Also, It is possible to extend PyADM1 linked to other models (similar to BSM2 approach) for plant wide simulation and development of digital twins.

Moreover, first principle mathematical models such as ADM1 are used in hybrid modeling for Industry 4.0 and Model Predictive Control (MPC) [14] as well as design, test, and simulation of Internet of Things (IoT) devices. Using Python in PyADM1 makes it easier to use in connection to common data science, machine learning, and web development libraries.

PyADM1 can serve as an educational tool [15, 16] to better demonstrate the anaerobic digestion process and different digester operational scenarios. Also, simulation of different scenarios is a low risk and low cost method to train biogas plant operators or evaluate process design by consultants.

Modeling and simulation of anaerobic digestion using PyADM1 can facilitate optimal design of experiments [17] for research about substrate utilisation and operational parameters. Plant designers and consultants use ADM1 to estimate plant performance [5], design parameters, and techno-economic studies. Models of wastewater treatment processes are important for process optimization and design [18], simulation of operational scenarios [15], and controller design [19].

## 5. Conclusions

PyADM1 was validated with the benchmark model BSM2 as a reliable Python implementation of ADM1 model for modeling and simulation of the anaerobic digestion process. The current version is capable of simulating steady-state operation of biogas reactors using ADM1 ODE/DAE forms. PyADM1 provides free transparent access to anaerobic digestion simulation for a broad Python user base as a low cost, reliable, and flexible solution for many applications, e.g. test and design of process control systems, biogas plant commissioning, design of experiments, operator training, and education.

Additional features such as dynamic simulation, data file read/write, application programming interface (API), and online graphical user interface will be added in the future versions. Linked to Python implementations of Activated Sludge Models (ASM) [20], PyADM1 can be a basis for open source wastewater treatment modeling and simulation software.

## Acknowledgements

This work was funded by Science Foundation of Ireland (SFI) Research Professorship Programme on Innovative Energy Technologies for Bioenergy, Biofuels and a Sustainable Irish Bioeconomy (*IETSBIO*^3^, award 15/RP/2763).

## Current executable software version

**Table 5:**
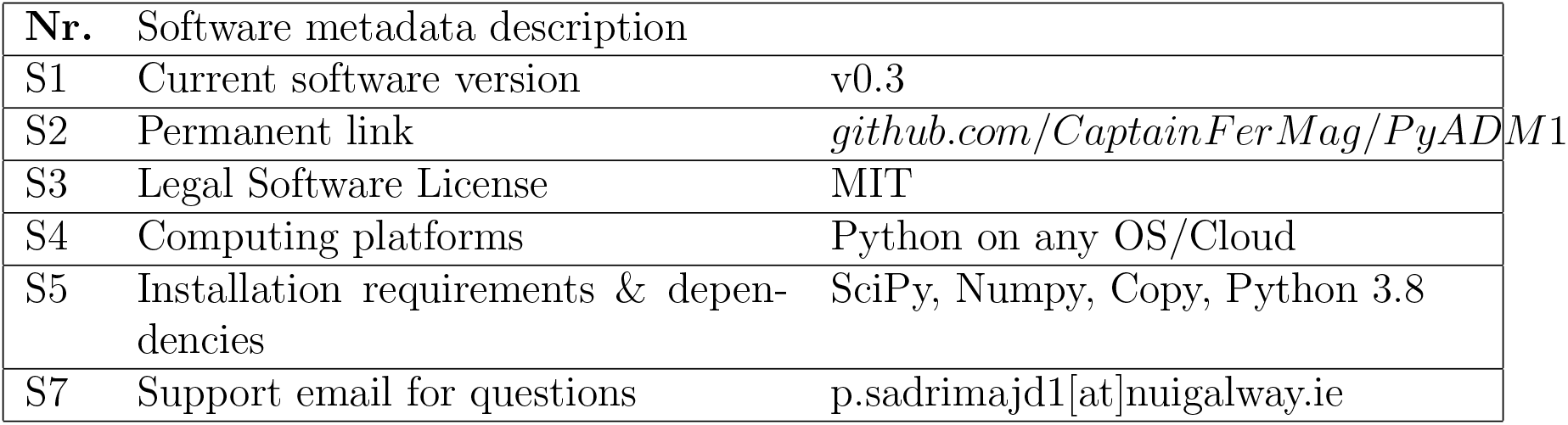
Software metadata

